# Impaired endogenous neurosteroid signaling contributes to behavioral deficits associated with chronic stress

**DOI:** 10.1101/2021.12.30.474579

**Authors:** Najah L. Walton, Pantelis Antonoudiou, Lea Barros, Alyssa DiLeo, Jenah Gabby, Samantha Howard, Rumzah Paracha, Edgardo J. Sánchez, Grant L. Weiss, Dong Kong, Jamie L. Maguire

## Abstract

Chronic stress is a major risk factor for psychiatric illnesses, including depression; however, the pathophysiological mechanisms whereby stress leads to mood disorders remain unclear. The recent FDA approval of antidepressants with novel mechanisms of action, like Zulresso®, a synthetic neuroactive steroid analog with molecular pharmacology similar to allopregnanolone, has spurred interest in new therapeutic targets and, potentially, novel pathophysiological mechanisms for depression. Allopregnanolone acts as a positive allosteric modulator of GABA_A_ receptors (GABA_A_RS), acting preferentially at δ subunit-containing receptors (δ-GABA_A_RS). Accumulating clinical and preclinical evidence supports the antidepressant effects of exogenous administration of allopregnanolone and allopregnanolone analogs; however, the role of endogenous neurosteroids in the pathophysiology of depression remains unknown. Here, we examine whether altered neurosteroid signaling may contribute to behavioral deficits following chronic unpredictable stress (CUS) in mice. We first identified reductions in expression of δ-GABA_A_Rs, the predominant site of action of 5a-reduced neuroactive steroids, following CUS. Additionally, utilizing LC-MS/MS we discovered a decrease in levels of allopregnanolone in the BLA, but not plasma of mice following CUS, an indication of impaired neurosteroid synthesis. CRISPR knockdown the rate-limiting enzymes involved in allopregnanolone synthesis, 5α-reductase type 1 and 2, in the BLA mimicked the behavioral deficits associated with CUS in mice. Furthermore, overexpression expression of 5α-reductase type 1 and 2 in the BLA improved behavioral outcomes. Collectively, this suggests chronic stress impairs endogenous neurosteroid signaling in the BLA which is sufficient to induce behavioral deficits similar to those observed following CUS. Further, these studies suggest that the therapeutic efficacy of allopregnanolone-based treatments may be due to their ability to directly target the underlying pathophysiology of mood disorders. Therefore, targeting endogenous neurosteroidogenesis may offer a novel therapeutic strategy for the treatment of mood disorders.

## Introduction

Recent advancements in the treatment of depressive disorders have successfully utilized synthetic neuroactive steroid analogs with molecular pharmacology similar to allopregnanolone which have been shown to exert both rapid and sustained antidepressant effects (Gunduz-Bruce, 2019; Meltzer-Brody et al., 2018). However, the mechanisms conferring antidepressant properties to this class of drugs is not entirely clear given that the predominant mechanism of action is as a positive allosteric modulator of GABA_A_ receptors (Belelli & Lambert, 2005; Chen et al., 2019; Mackenzie & Maguire, 2013) a target which has not yet been extensively linked to the underlying pathophysiology of depression, although our work has implicated these receptors in the pathophysiology of postpartum depression (Maguire and Mody, 2008; Walton and Maguire, 2019).

GABA_A_ receptors are heteropentameric ligand-gated ion channels which mediate inhibitory neurotransmitter signaling in the brain. GABA_A_ receptor properties, including pharmacological sensitivity, are dictated by subunit composition, in which γ2-subunit containing GABA_A_ receptors are sensitive to benzodiazepines and extrasynaptic δ-subunit containing GABA_A_ receptors (δ-GABA_A_RS) confer sensitivity to neurosteroids (Belelli et al. 2002). Interestingly, we recently demonstrated that neurosteroid-sensitive δ-GABA_A_RS are uniquely expressed on parvalbumin (PV) interneurons in the basolateral amygdala (BLA) and orchestrate network states that govern behavioral states (Antonoudiou et al., 2021; Davis et al., 2017; Ozawa et al., 2020). As such, neurosteroid-mediated potentiation of these receptors has been demonstrated to modulate BLA network and behavioral states (Antonoudiou et al., 2021). Further, exogenous neurosteroid treatment prevents or restores the network and behavioral deficits associated with chronic stress (Antonoudiou et al., 2021). Thus, we propose that perturbations in neurosteroid signaling in the BLA may contribute to network and behavioral state changes.

Deficits in neurosteroids have been demonstrated following chronic stress (Dong et al., 2001; Matsumoto et al., 2005; Serra et al., 2006; Serra et al., 2000) which we propose may contribute to the behavioral deficits associated with chronic stress. Further, there is a reduction in the expression of the rate limiting enzymes involved in endogenous neurosteroidogenesis, 5α-reductase, following chronic stress (Agís-Balboa et al., 2007; Dong et al., 2001). Given that chronic stress is a major risk factor for psychiatric illnesses (McEwen & Akil, 2020; McEwen, 2017; McEwen, 1998), these data suggest that deficits in endogenous neurosteroidogenesis may play a role in the underlying pathophysiological mechanisms of psychiatric disorders (van Broekhoven and Verkes, 2003; Zorumski et al., 2013).

Here, we directly examine the contribution of endogenous neurosteroid signaling in the BLA in mediating changes in behavioral states associated with chronic stress. These studies build upon the well-established evidence that chronic stress is a risk factor for psychiatric disorders (McEwen, 2017; McEwen & Akil, 2020) and can be utilized in rodents to alter behavioral outcomes (Juster et al. 2010; Willner, 2017). In these studies, we demonstrate that chronic unpredictable stress (CUS) induces changes in the capacity for endogenous neurosteroid signaling in the BLA, evident from a decrease in allopregnanolone levels in the BLA but not circulating plasma, decreased expression of the neurosteroidogenic enzymes, 5α-reductase 1 and 2 (5α1/2), and their main site of action, δ-GABA_A_RS in the BLA. CRISPR-mediated knockdown of the genes encoding for 5α-reductase 1 and 2, Srd5a1 and Srd5a2, respectively, is sufficient to mimic the impact of CUS on behavioral state changes. Importantly, increasing the capacity for endogenous neurosteroidogenesis by overexpressing 5α1/2 in the BLA is sufficient to improve behavioral states. These findings implicate endogenous neurosteroid signaling in the BLA in the underlying pathophysiology of mood disorders and provides a potential mechanistic underpinning for the antidepressant effects of 5α-reduced neurosteroids.

## Methods and Materials

### Animals

Adult (3-month-old) male (Stock No. 000664) and constitutively expressing Cas9 mice (Stock No. 024858) were purchased directly from The Jackson Laboratory. Mice were housed at Tufts University School of Medicine’s Division of Laboratory Animal Medicine facility in clear plastic cages (4 mice per cage), in a temperature and humidity-controlled facility, on a 12 h light/dark cycle (lights on at 0700 h), with ad libitum access to food and water. All procedures were approved by the Tufts University Institutional Animal Care and Use Committee.

### Chronic Unpredictable Stress (CUS)

Adult male C57BL/6J mice underwent a three-week CUS protocol consisting of four consecutive days of alternating stressors only during the dark period, including cage tilt, restricted cage, food and water restriction, soiled cage, and unstable cage, followed by an acute stressor on the fifth day (restraint stress, 2-hours of cold stress exposure or tail suspension) (Supplemental figure 1). Control mice were gently handled by the experimenter twice per week.

### Viral Production

Cre-dependent adeno-associated virus (AAV) viral vectors were constructed as previously described (Xu et al. 2018) and packaged at the Boston Children’s Hospital Viral Core facility. In brief, AAVs were constructed and cloned on a pAAV-pEF1α-FLEX- mCherry-WPRE-pA plasmid. sgRNAs were first designed utilizing online CRISPR tools (http://crispr.mit.edu/ and http://chopchop.cbu.uib.no/). The pU6-sgRNA-scaffold cassette in each guide was designed with an in-house Snap-Ligation kit, to facilitate expression of sgRNAs in each AAV vector. The sgRNA-scaffold cassette was subsequently cloned into the Mlul site of the plasmid backbone. The following AAV coat serotypes and viral titers (viral molecules/mL) were used: pAAV-ef1a-U6-srd5a1gRNA-DIO-mcherry (2/8, 7.75E+12) and pAAV-ef1a-U6-srd5a2gRNA-DIO-mcherry (2.8, 1.04E+13). Viral aliquots were stored at −80°C before stereotaxic injection.

### Stereotaxic surgery

Mice were anesthetized with an intraperitoneal injection of 100 mg/kg ketamine and 10 mg/kg xylazine mixture. Before beginning the surgical procedure, mice were administered sustained release buprenorphine (0.5mg/kg, subcutaneous injection). Burr holes in the skull were created to deliver 250nl of guide RNAs (sgSrd5a1 and sgSrd5a2) bilaterally into the BLA (AP −1.5mm; ML +/- 3.3mm; DV - 5.1mm) of constitutively expressing Cas9 mice using a 33-gauge Hamilton syringe at an infusion rate of 100nL/minute. Lentiviral overexpression of Srd5a1 (Origene Cat No. MR203282) and Srd5a2 (Origene Cat No. MR223814) in the BLA was performed in the same manner using C57BL/6J mice. Post Hoc analysis was performed to confirm appropriate targeting and expression. qRT-PCR was performed to validate knockdown or overexpression of the enzymes.

### Quantitative Real-Time PCR (qRT-PCR)

qRT-PCR was performed as previously described (Melón et al., 2018) to quantify mRNA levels of *Srd5a1* and *Srd5a2* in control and CUS male and female mice. In brief, mice were anesthetized with isoflurane, decapitated, the brain rapidly extracted, flash frozen in liquid nitrogen, and stored at −80°C until further processing. Tissue punches from the BLA were collected using a 0.5 mm sterile biopsy needle and subsequently homogenized in 25μL of TRIzol™. Chloroform (15μl) was then added, samples were centrifuged, and the supernatant was transferred to a sterile tube containing 12.5μL of cold isopropanol and 2μL of glycogen. Samples were then pelleted and rinsed in 75% ethanol, air dried, and the resulting RNA product was heated at 60°C in 1mM of sodium citrate made up in RNAsecure. RNA integrity and concentration were measured using a Thermo Scientific NanoDrop 1000. SuperScript III Platinum Sybr Green (one step qRT PCR kit) (Invitrogen Lot No. 2028735) was used for SYBR-based qRT-PCR along with 100ng of template RNA and primers listed in Table 2. A Stratagene Mx3000P (Agilent Genomics, Santa Clara, CA, USA) was used to perform the qRT-PCR using the following thermocycling parameters: 1 cycle at 50°C for 3 minutes, 1 cycle at 95°C for 5 minutes, 40 cycles of 15 seconds at 95°C followed by 40 seconds at 60°C, 1 cycle at 40°C for 1 minute, 1 cycle at 95°C for 1 minute followed by 30 seconds at 55°C and 30 seconds at 95°C. All samples were run in duplicate with an average CT value normalized to β-actin and relative transcript levels were calculated in accordance with the 2^-ΔΔCT^ method (Melón et al., 2018).

**Table 1.**
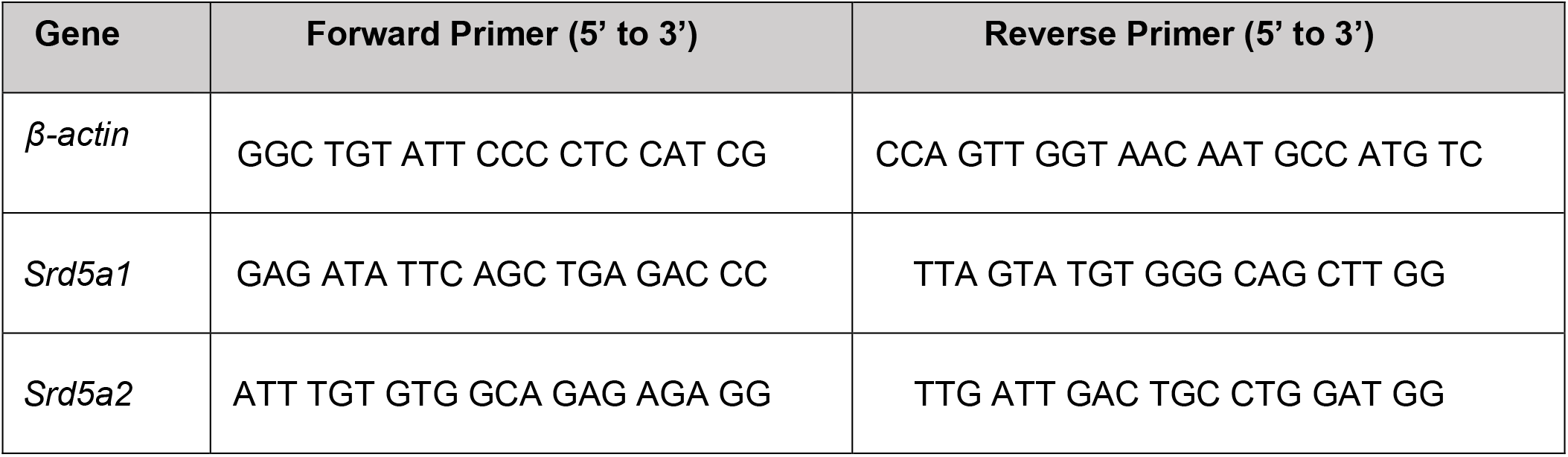
List of IDT custom primers used for qRT-PCR. *Srd5a1* and *Srd5a2* mRNA levels were normalized to *β-actin*.

### Animal Behavior

#### Elevated Plus Maze

Avoidance behaviors were assessed using the elevated plus maze as previously described (Melón et al., 2018; Antonoudiou, 2021). In brief, mice were individually placed in the center of the apparatus randomly facing either a closed or an open arm before beginning the 10-minute trial. The plus maze is elevated 75cm in height, with 4 arms (38cm x 6.5cm) and each of the two closed arms have walls 10cm in height. At the completion of testing the total distance traveled, time spent in open and closed arms, and the number of entries into the closed and open arms were analyzed using automated Motor Monitor software (Hamilton-Kinder).

#### Open Field

Locomotor and avoidance behaviors were assessed in a brightly illuminated open field as previously described (Antonoudiou, 2021). Open field testing was performed by placing mice in the center of the chamber in alternating directions per subject and activity was recorded over a 10-minute period using automated Motor Monitor software (Hamilton-Kinder). The open field chamber consists of a 40 cm x 40 cm plexiglass open field chamber surrounded by a photobeam frame with equally spaced photocells to detect the total distance traveled (cm), time spent in the periphery and center of the chamber (sec), rearing behavior, and overall number of basic movements made during the test.

#### Light/Dark Box

Avoidance behavior was also assessed in the light/dark box as previously described (Melón et al., 2018; Antonoudiou, 2021). Mice were placed in an enclosed dark compartment of the two-chamber apparatus. Upon initiation of the 10-minute trial the barrier between the two chambers was removed to allow mice free access to both light and dark chambers. The apparatus was enclosed by a 22 x 43 cm photobeam frame with equally spaced photocells. Distance traveled, the amount of time spent in the light compartment, and the number of entries into the light compartment were analyzed using automated Motor Monitor software (Hamilton-Kinder).

#### Tail Suspension

Learned helplessness was assessed using the tail suspension test as previously described (Lee et al., 2014; Antonoudiou, 2021). In brief, mice were suspended by their tail on a bar 36cm above a platform for 6 minutes. Trials were video recorded and automatically scored using an unbiased in-house Mobility App (https://github.com/researchgrant/mobility-mapper.git). Limb movement (nose, paws, and tail) were calculated from positional data automatically detected using DeepLabCut (Mathis 2018). A neural network was then trained using a Long Short-Term Memory recurrent neural network architecture in the TensorFlow toolbox for Python to predict the mobility state of the animals based on videos scored by several different investigators. The mobility network had a frame-by frame accuracy of over 90% and overall mobility time correlated to manual scored data, r^2^= 0.91. Mice were excluded from data analysis if they were able to free themselves from suspension or climbed their tail during any point in the trial.

### Statistical Analysis

All statistical analysis was performed using GraphPad Prism 9 software, MATLAB (MathWorks, 2020), and Python (Python Software Foundation). Two-tailed paired t-tests were used to analyze within animal conditions, unpaired t-tests were used to analyze between two experimental groups. Values reported herein are as mean ± SEM. P-value denotation is as follows: < than 0.05 were considered statistically significant; p ≥0.05=n.s. p<0.05=*, p<0.01=**, p<0.001=***, p<0.0001=****.

## Results

### Behavioral Aberrations Following CUS

Given that chronic stress is a well-established risk factor for the development of psychiatric disorders, we utilized a chronic unpredictable stress rodent model to examine the behavioral sequelae of chronic stress. Similar to our previous studies (Antonoudiou, 2021), mice subjected to CUS displayed increased avoidance behaviors and learned helplessness compared to naive animals. Following CUS, mice exhibit a shorter latency to immobility (control: 73.50 +/- 7.418 s; CUS: 35.91 +/5.947 s) (Figure 1A; control n=10, CUS n=11, p=0.0008 unpaired t-test) as well as a greater overall time immobile (control: 180 +/- 13.76 s; CUS: 233.2 +/- 7.754 s) in the tail suspension test (Figure 1A; p=0.0027 unpaired t-test). Similarly, mice subjected to CUS exhibit a decreased latency to immobility (control: 67.20 +/- 4.790 s; CUS: 11 +/- 3.225 s) and an increased total time immobile (control: 149 +/16.11 s; CUS: 226.4 +/- 7.633 s) in the forced swim test (Figure 1B; latency: control n=5, CUS n=5, p=<0.0001, total time: p=0.0055; unpaired t-test). Mice that underwent CUS also displayed increased avoidance behaviors, evident from a significant decrease in the amount of total time spent in the center of the open field compared to control mice (control: 70.72 +/- 13.48 s; CUS: 32.70 +/- 2.922 s) (Figure 1C; control n=5, CUS n=8, p=0.0054 unpaired t-test). However, there were no changes in overall locomotor behavior (control: 2063 +/- 245.5 beam breaks; CUS: 2337 +/- 140.4 beam breaks) (Figure 1C; p=0.3372 unpaired t-test). Thus, CUS induces robust changes in behavior, making this model and these outcome measures useful for investigating the mechanisms whereby stress alters behavioral outcomes.

**Figure 1.**
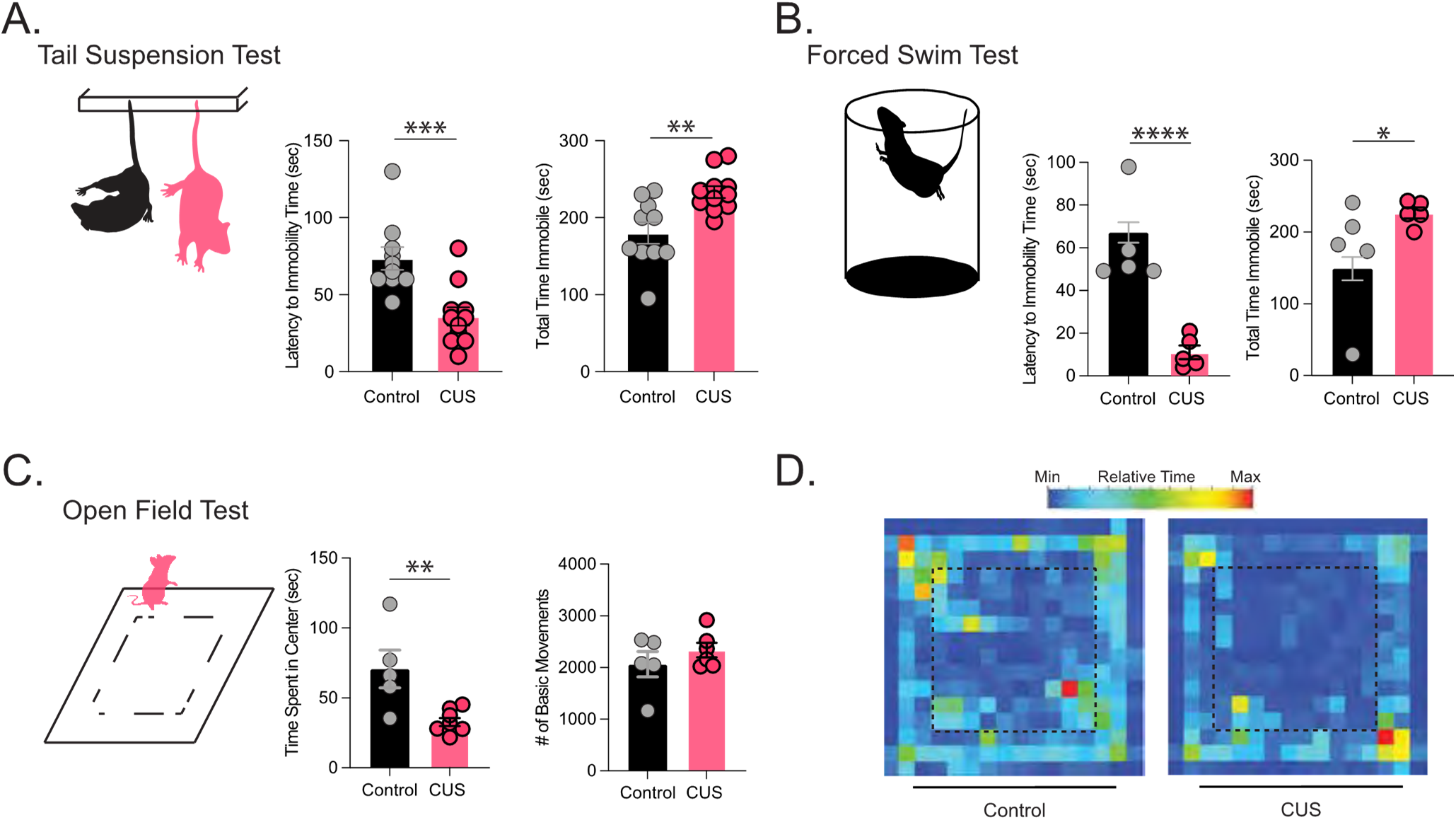
Chronic unpredictable stress induces behavioral deficits. CUS increases learned helplessness, decreasing the latency to immobility time and total time spent immobile during tail suspension control n=10, CUS n=11(A) and forced swim tests control n=5, CUS n=5 (B). CUS also increases avoidance behaviors, decreasing the total time spent in the center of the open field without altering overall locomotor behavior control n=5, CUS n=8(C), which can be appreciated in a representative heat plot of open field behavior in control and CUS mice (D). * Denotes p < 0.05 using an unpaired Student’s t-test.

### Impaired Neurosteroidogenesis in the BLA Following CUS

While exogenous treatment with neurosteroid analogs has demonstrated clinical effectiveness in the treatment of depressive disorders, there remains questions as to how these treatments exert their effects and whether they target the underlying pathophysiology of depressive disorders. To investigate this gap in knowledge we sought to determine whether endogenous neurosteroidogenesis is impacted by chronic stress. Allopregnanolone levels were measured in plasma, whole brain, and BLA samples from control and CUS mice using LC-MS. Plasma levels of allopregnanolone were not significantly different between control and CUS mice (control: 1.63 +/- 0.800324817 ng/ml; CUS: 0.59 +/- 0.08417 ng/ml) (Figure 2A; control n=11, CUS n=13, p=0.087747 unpaired t test). Similarly, there was no significant difference in the total brain levels of allopregnanolone (control: 3.23 +/- 1.14550076 ng/g; CUS: 1.88 +/- 0.761861809 ng/g) (Figure 2B; control n=6, CUS n=4, p= 0.873461 unpaired t test). However, we observed a decrease in allopregnanolone levels measured in the BLA following CUS compared to controls (control: 3.8 +/- 1.065780826 ng/g; CUS: 1.77 +/- 0.311052311 ng/g) (Figure 2C; control n=4, CUS n=5, p= 0.041426 unpaired t test). These data suggest that there is an impairment in endogenous neurosteroid synthesis following CUS and that these deficits are likely brain region-specific.

**Figure 2.**
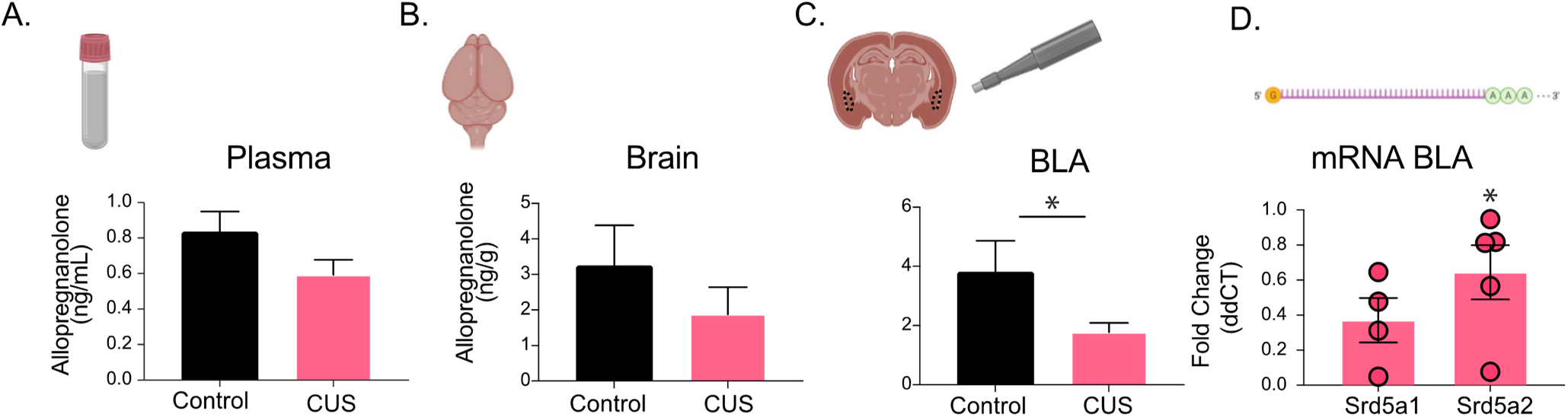
Impaired endogenous allopregnanolone in the BLA following CUS. LC-MS measurement of allopregnanolone levels from control and CUS (A) plasma, control n=11, CUS n=13 (B) brain, control n=6, CUS n=4 (C) BLA samples, control n=4, CUS n=5 (D) qRT-PCR measurement of BLA mRNA levels of Srd5a1 and Srd5a2 normalized to β-actin and control levels, control n=16, CUS n=16. * denotes p < 0.05 using an unpaired Student’s t-test.

Considering the changes in endogenous neurosteroid levels measured in the BLA, we assessed the levels of 5α-reductase 1 and 2 expression in the BLA given that these are the rate-limiting enzymes for allopregnanolone synthesis. qRT-PCR revealed a decrease in BLA transcript levels of *Srd5a1* and *Srd5a2*, which encode for 5α-reductase 1 and 2, respectively, following CUS (Srd5a1: 0.3710 +/- 0.1273 fold change; Srd5a2: 0.6443 +/- 0.1548 fold change) (Figure 2D; control n=16 pooled BLA samples from 4 mice per run, CUS n=16 pooled BLA samples from 4 mice per run, 4 runs Srd5a1 p=0.0617; 5 runs Srd5a2 p=0.0141 one sample t test). These data suggest that endogenous neurosteroidogenesis is impaired in the BLA following CUS, which we proposed may contribute to the behavioral deficits.

### Impaired Neurosteroidogenesis in the BLA is Sufficient to Induce Behavioral Deficits

Given our evidence that CUS induces brain region-specific deficits in endogenous neurosteroidogenesis, we next sought to directly investigate whether the behavioral consequences of CUS involve impaired neurosteroidogenesis in the BLA. Utilizing constitutive Cas9 expressing mice we knocked down both 5α-reductase 1 and 2 (5α1/2) in the BLA using sgRNAs targeting the genes encoding these enzymes, *Srd5a1* and *Srd5a2* respectively. Both enzymes were targeted and constitutive Cas9 was utilized to target all cell types given the current lack of information about the function of these enzymes and cell types involved in neurosteroidogenesis in the BLA. This approach significantly reduced the transcript levels of both 5α-reductase 1 and 2 (Srd5a1: 0.4704 +/- 0.07069 fold change; Srd5a2: 0.7219 +/- 0.06515 fold change) in the BLA (Figure 3B control n=15, sgSrd5a1-2 n=8; Srd5a1 p=0.0003; Srd5a2 p=0.0001). Mice with reduced 5α-reductase expression exhibited increased avoidance behaviors, spending significantly less time in the light compartment of the light/dark box (control: 286.7 +/- 21.69 s; 5α1/2: 233 +/- 13.46 s) (Figure 3C control n=15, sgSrd5a1-2 n=8, p=0.0480 unpaired t test). Mice with a knockdown of 5α1/2 expression also displayed increased learned helplessness, evident from a shorter latency to immobility (control: 71.92 +/- 3.736 s; sgSrd5a1-2: 31.67+/- 5.652 s) (Figure 3D p=<0.0001 unpaired t test) and an increase in the total time immobile (control: 173.2 +/- 10.28 s; sgSrd5a1-2: 237.2 +/- 13.05 s) (Figure 3D p=0.0013) in the tail suspension test. Collectively, these results suggest that impaired neurosteroid signaling in the BLA as a result of knockdown of key neurosteroidogenic enzymes 5α1/2, contributes to behavioral abnormalities which mimic those observed following chronic stress. Further, these data demonstrate that endogenous 5α-reduced neurosteroids exert a tonic influence over behavioral states.

**Figure 3.**
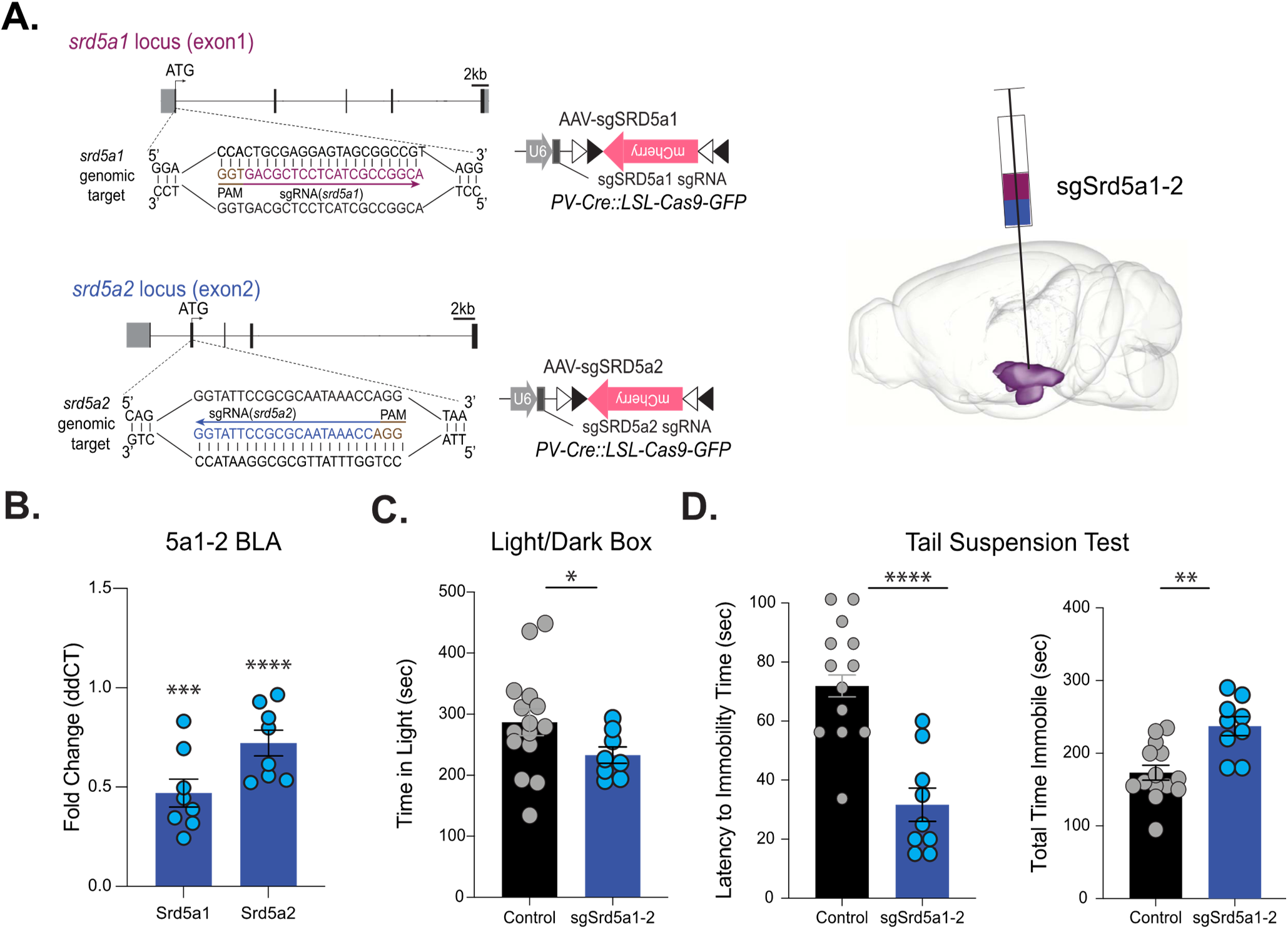
Knockdown of Srd5a1 and Srd5a2 in the BLA induces behavioral deficits. (A) left diagram of sgRNA construct for knockdown of Srd5a1 and Srd5a2 in constitutive Cas9 mice. Right schematic of targeting for sgRNA in the BLA. (B) qRT-PCR measurement of Srd5a1 and Srd5a2 mRNA levels in sgSrd5a1-2 mice. Results are normalized to b-actin levels and expression in controls. Knockdown of 5α-reductase 1 and 2 increases avoidance behaviors, decreasing the total amount of time spent in the light chamber of the light/dark box (C). Knockdown of 5α-reductase 1 and 2 increases learned helplessness, decreasing the latency to immobility time, and increasing the total time spent immobile during the tail suspension test (D). Control n=15, sgSrd5a1-2 n=8 per experimental group. * denotes p < 0.05 using an unpaired Student’s t-test.

### Enhanced Neurosteroidogenesis in the BLA is Sufficient to Improve Behavioral States

Given that a reduction in 5α1/2 expression in the BLA is sufficient to increase avoidance behaviors and learned helplessness, we next sought to investigate how rescue of 5α1/2 in the BLA may impact behavioral outcomes. To do so, we utilized lentiviral constructs to overexpress 5α1/2 in the BLA and assessed behaviors impacted by chronic stress, including avoidance behaviors, and learned helplessness. This approach significantly increased both 5α1 and 2 transcript levels in the BLA (Srd5a1: 1.950 +/- 0.3382 fold change; Srd5a2: 1.549 +/- 0.1512 fold change) (Figure 4A control n=7, LV Srd5a1-2 n=4; Srd5a1 p=0.0104; Srd5a2 p=0.0020). Mice with 5α1/2 overexpressed in the BLA (LV-Srd5a1-2 mice) spent a greater amount of time (control: 53.97 +/- 7.456 s; LV-Srd5a1-2: 76.75 +/- 7.845 s) and traveled further in the center of the open field (control: 529.4 +/- 83.81 cm; LV-Srd5a1-2: 994.8 +/- 56.96 cm) (Figure 4A; control n=14, LV Srd5a1-2 n=10, p=*0.0476, ***0.0002 unpaired t test). Additionally, LV-Srd5a1-2 mice completed more entries into the center of the open field as compared to controls (control: 23.46 +/- 4.794 entries; LV-Srd5a1-2: 46.20 +/- 3.169 entries) (Figure 4A; p=0.0008 unpaired t test) and displayed an increase in exploratory behavior in this task (control: 2192 +/- 183.4 beam breaks; LV-Srd5a1-2: 2919 +/- 147.2 beam breaks) (Figure 4A; p=0.0053 unpaired t test). In the elevated plus maze LV-Srd5a1-2 mice made more entries into the open arm (control: 31.03 +/- 6.026 entries; LV-Srd5a1-2: 48.30 +/- 10.16 entries) (Figure 4B p=0.0001 unpaired t test) indicating a decrease in avoidance behaviors in these mice. LV-Srd5a1-2 mice also exhibit decreased learned helplessness behaviors, exhibiting an increased latency to immobility (control: 71.92 +/- 3.726 s; LV-Srd5a1-2: 98.89 +/- 5.879 s) (Figure 4C; p=0.00017 unpaired t test) as well as a decrease in total time immobile (control: 173.2 +/- 10.28 s; LV-Srd5a1-2: 122.5 +/- 12.68 s) in the tail suspension test (Figure 4C; p=0.0058 unpaired t test). These data demonstrate that enhancing the synthesis of 5α-reduced neurosteroids improves behavioral states.

**Figure 4.**
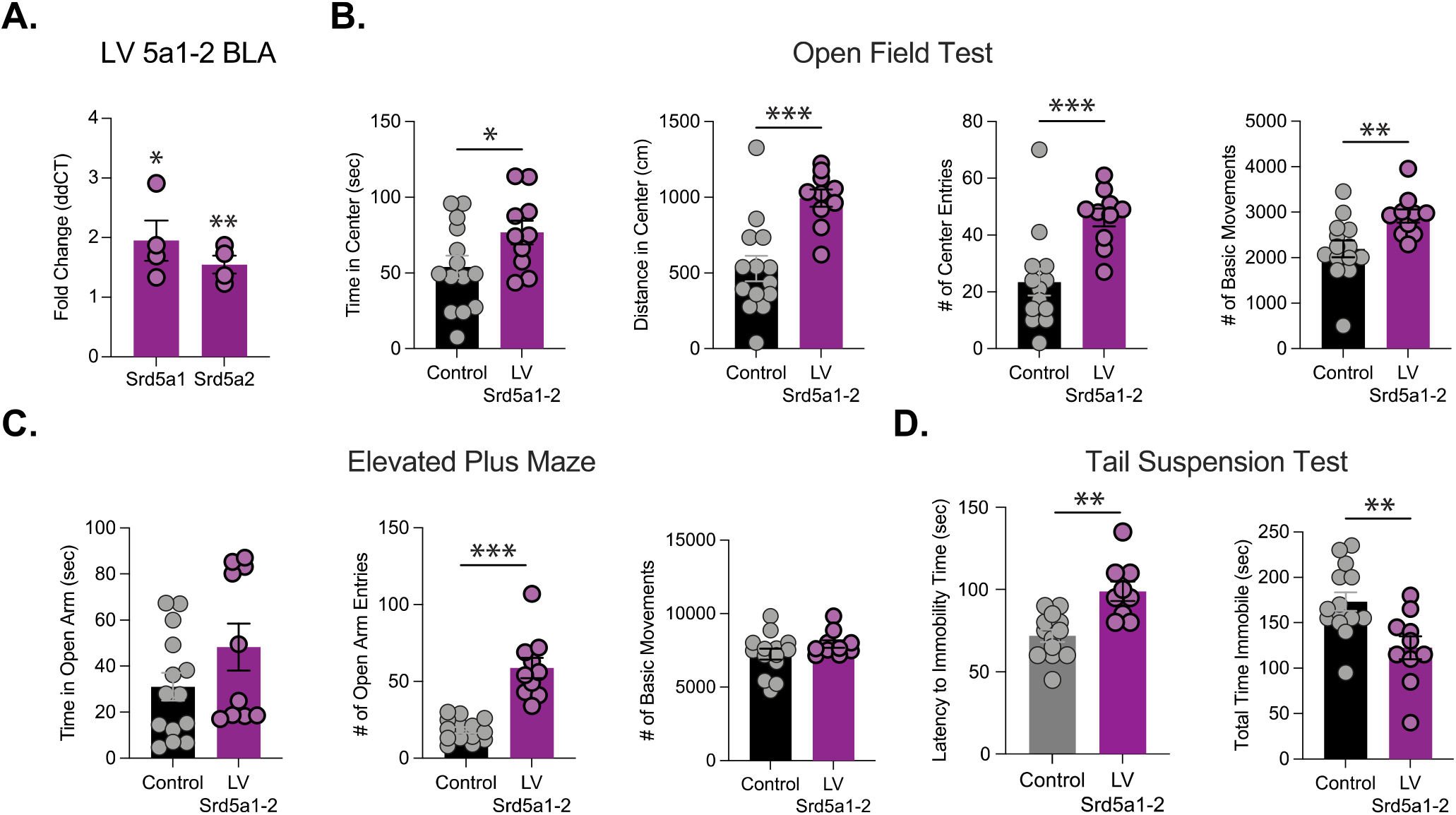
Overexpression of Srd5a1 and Srd5a2 improves behavioral states. qRT-PCR measurement of Srd5a1 and Srd5a2 mRNA levels in LV Srd5a1-2 mice. Results are normalized to b-actin levels and expression in controls, control n=7, LV Srd5a1-2 n=4(A). Overexpression of 5α-reductase 1 and 2 decreases avoidance behaviors, increasing the time in the center, distance traveled in the center, and the number of entries into the center of the open field test (B). Overexpression of 5α-reductase 1 and 2 increases the number of open arm entries but not the total time in the open arm (C). Overexpression of 5α-reductase 1 and 2 decreases learned helplessness, increasing the latency to immobility and the total time spent immobile (D). Control n=14, LV Srd5a1-2 n=10 per experimental group. * denotes p < 0.05 using an unpaired Student’s t-test.

## Discussion

Here we demonstrate a novel mechanism through which chronic stress alters behavioral states, involving deficits in endogenous neurosteroid signaling in the BLA (Figure 5). Previously, our laboratory demonstrated deficits in network and behavioral states following CUS which were prevented or reversed with exogenous allopregnanolone treatment (Antonoudiou et al. 2021). As a mechanistic follow up to the previous study, here we investigated whether deficits in endogenous neurosteroid signaling may contribute to the underlying pathophysiology of depression, utilizing a chronic stress paradigm. These novel findings further our understanding of the mechanisms through which chronic stress alters mood and how and why exogenous 5α-reduced neurosteroids exert antidepressant effects.

**Figure 5.**
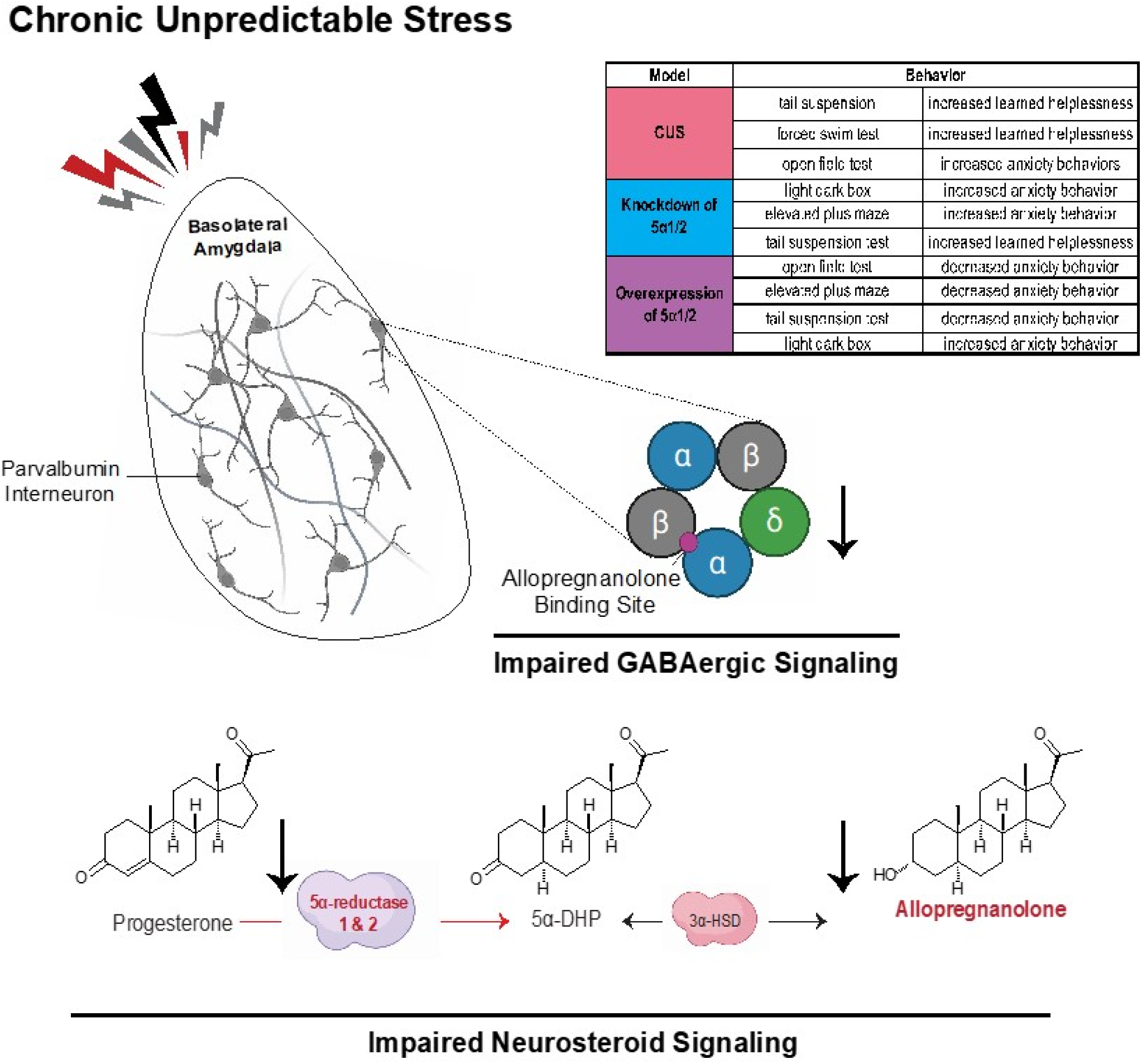
Schematic overview of major findings. Depiction of the impact chronic unpredictable stress has upon the BLA. Findings support impaired endogenous neurosteroid signaling following CUS, evident from a downregulation of 5α1-2 and decreased allopregnanolone levels in the BLA. Knockdown of 5α-reductase 1 and 2 mimics the behavioral deficits associated with chronic stress. Conversely, overexpression of 5α-reductase 1 and 2 improves behavioral states. Table in the top right summarizes the major findings in each experimental model used in this paper.

In an effort to understand the mechanisms mediating transitions between network and behavioral states, we previously identified PV interneurons in the BLA as critical mediators in generating oscillations that govern behavioral state changes and also determined that they contain a high density of neurosteroid-sensitive, δ-GABA_A_Rs (Antonoudiou et al., 2021). Thus, it is likely that neurosteroid actions on δ-GABA_A_Rs on PV interneurons in the BLA mediates the ability of allopregnanolone to shift network and behavioral states. However, it was previously unclear whether exogenous allopregnanolone treatment exerted its antidepressant effects by targeting the underlying pathophysiological mechanism. Here we demonstrate deficits in endogenous neurosteroid signaling in the BLA following CUS (Figure 5), suggesting that the impact of exogenous allopregnanolone on network and behavioral states may in fact be targeting the underlying pathophysiological mechanism. Further, we demonstrate that inducing deficits in endogenous neurosteroid signaling in the BLA, through a knockdown of 5α-reductase 1 and 2 in the BLA, is sufficient to recapitulate the behavioral deficits observed in mice following CUS (Figure 3 and Figure 6). These findings are consistent with the observed reduction in the expression of 5α-reductase (Agís-Balboa et al., 2007; Dong et al., 2001) and allopregnanolone levels (Dong et al., 2001; Matsumoto et al., 2005; Serra et al., 2006; Serra et al., 2000) previously observed in chronic stress models and implicated in the mechanisms contributing to psychiatric disorders (van Broekhoven and Verkes, 2003; Zorumski et al., 2013).

Prior work investigating the role of neurosteroids in affective disorders emphasize the antidepressant and anxiolytic effects of these compounds (Zorumski et al., 2013). The ability of neurosteroids to influence mood have been suggested to involve numerous mechanisms, including the well-established action of 5α-reduced neurosteroids as positive allosteric modulators of GABA_A_RS, the ability to regulate inflammation, and/or neuroprotective effects (Walton and Maguire, 2019; Zorumski, 2019). The current manuscript and other recent work from our laboratory suggest that the antidepressant mechanism of allopregnanolone and synthetic neuroactive steroid analogs with molecular pharmacology similar to allopregnanolone may involve the ability of these compounds to modulate BLA network states, influencing affective switching and contributing to the persistent antidepressant effects of these compounds (Antonoudiou et al. 2021; Schiller et al. 2014).

The ability of neurosteroids to mediate transitions between BLA network and behavioral states likely involves the actions of these compounds on δ-GABA_A_RS expressed on PV interneurons in the BLA (Antonoudiou et al. 2021) and the ability of tonic inhibition on interneurons to synchronize network activity (Pavlov et al. 2014). We propose that impaired neurosteroid-mediated enhancement of tonic GABAergic signaling on PV interneurons in the BLA following CUS impairs the ability to effectively synchronize network states and, thereby, behavioral states (Figure 5).

Most studies investigating the influence of neurosteroids on mood have relied on the exogenous administration of neurosteroids. The current study extends our knowledge by providing insight into the role of endogenous neurosteroid signaling on behavioral states. Currently, very little is known about the function of endogenous neurosteroidogenesis. In the clinical population, treatment with finasteride has been shown to lead to mood disorders, including anxiety and depression, collectively referred to as post-finasteride syndrome (PFS), the pathophysiology of which is thought to involve deficits in endogenous neurosteroid signaling (Melcangi et al., 2013). In animal models, treatment with allopregnanolone exerts anxiolytic and antidepressant effects and treatment with finasteride blocks the anxiolytic and antidepressant effects of progesterone (for review see (Finn et al., 2006)). Collectively, these data suggest that endogenous neurosteroids are capable of impacting mood. Here we directly demonstrate deficits in endogenous neurosteroid signaling associated with a major risk factor for psychiatric illnesses, chronic stress. The current study demonstrates that interfering with endogenous neurosteroid signaling is capable of inducing deficits in behavioral states, suggesting that endogenous neurosteroids have a tonic influence over affective states. Further, we demonstrate that enhancing endogenous neurosteroidogenesis is capable of improving behavioral states, suggesting the possibility that targeting endogenous neurosteroidogenesis may represent a novel therapeutic approach for the treatment of mood disorders (Porcu et al., 2016; Pinna, 2014). Further studies are required to investigate the therapeutic potential of targeting endogenous neurosteroidogenesis for the treatment of mood disorders.

## Acknowledgements and Disclosures

The authors would like to thank Boston Children’s Hospital viral vector core for packaging the guideRNAs used in this manuscript as well as PharmaCadence for performing LC-MS analytical support. JLM, NLW, PA, AD, and RP were supported by R01AA026256, R01NS105628, R01NS102937, R01MH128235, P50MH122379, and a sponsored research agreement with SAGE Therapeutics. SH was supported by T32DK124170. DK was supported by R01NS107315, R01DK108797, R21HD098056, P30DK046200. JLM serves as a member of the Scientific Advisory Board for SAGE Therapeutics, Inc. All other authors report no potential biomedical financial interests or conflicts of interest.

## Notes

### Competing Interest Statement

Jamie Maguire serves on the Scientific Advisory Board for SAGE Therapeutics

