## Supplemental Materials for "Impaired endogenous neurosteroid signaling contributes to behavioral deficits associated with chronic stress"

**Supplemental Material**

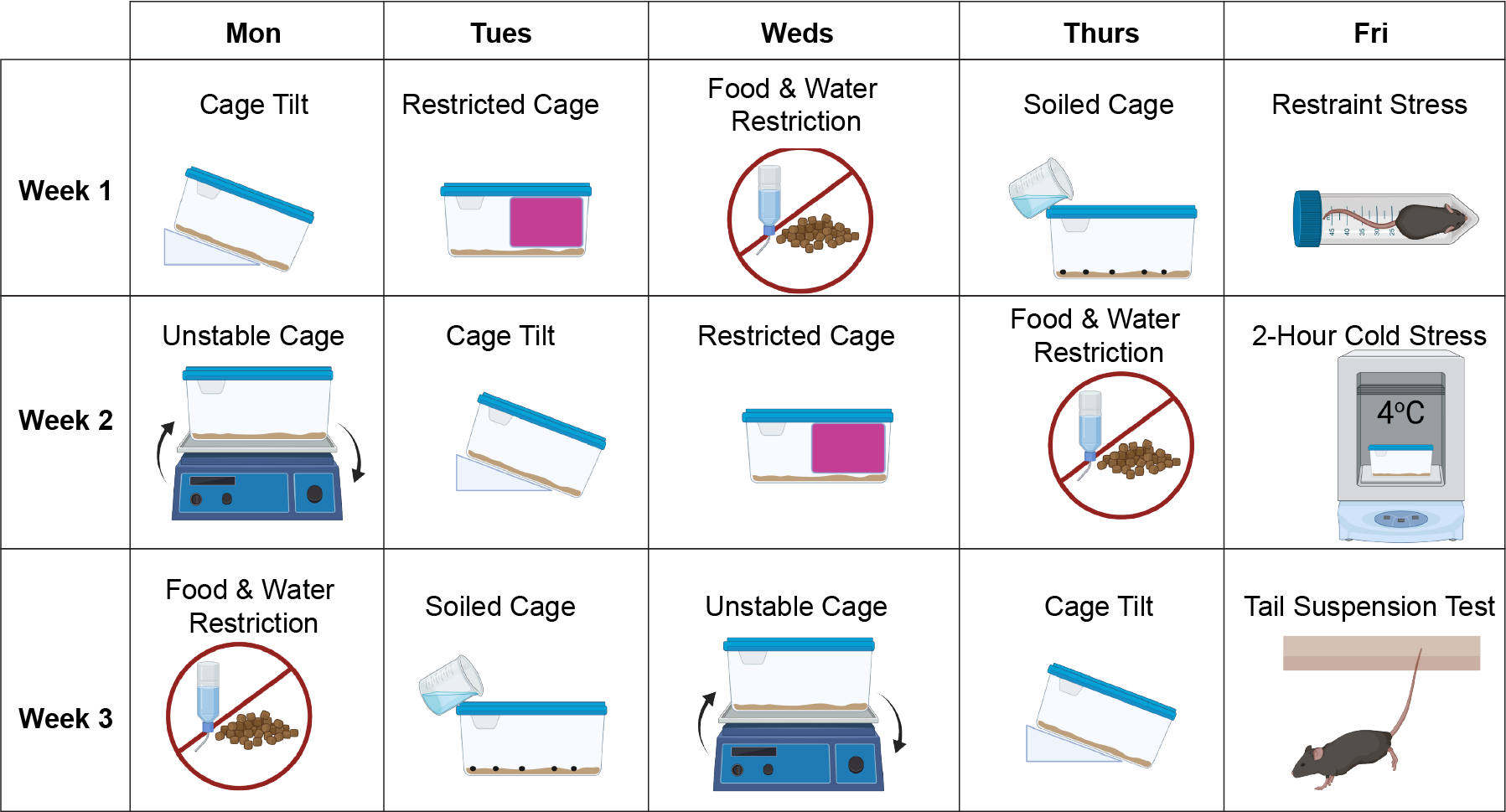

**Supplemental Figure 1. Timeline of CUS protocol.** Schematic overview of the CUS protocol consisting of 4 days of alternating overnight stressor (cage tilt, restricted cage, food and water restriction, soiled cage, and unstable cage) followed by a fifth day of an acute stressor (restraint stress, 2-hours of cold stress exposure, and tail suspension). Stressor and start times were randomized. This protocol was repeated for 3 weeks as shown.
